# An Adjustable Offloading Ankle-Foot Orthosis: Design and Proof-of-Concept Biomechanical Verification

**DOI:** 10.64898/2026.05.17.725313

**Authors:** Eshraq Saffuri, Lee Jordan, Dana Solav

## Abstract

Various ankle-foot conditions (e.g., fractures, diabetic foot ulcers, and post-surgical recovery) require periods of complete non-weightbearing followed by gradually increasing partial loadings. However, existing assistive devices often provide inconsistent or uncomfortable offloading during gait. Additionally, prolonged proximal leg offloading can contribute to muscle atrophy, reduced bone density, and overuse of other body segments. We present a novel offloading ankle-foot orthosis (OLAFO) designed to overcome these limitations. The OLAFO features a patient-specific load-bearing shank brace, designed through a digital workflow and fabricated from a 3D-printed core reinforced with carbon-fiber composite lamination. Interlocking serrated side struts, adjustable in 2 mm increments, modulate load sharing between the shank and plantar surfaces. Furthermore, the OLAFO incorporates contact plates with a rocker profile informed by roll-over-shape measurements to support forward progression and gait symmetry. Proof-of-concept biomechanical verification in one able-bodied participant evaluated complete offloading, five partial-loading levels, and normal gait using a pressure walkway to compute vertical ground reaction forces and impulses. In complete offloading, the affected foot generated no contact pressures. Across partial-loading levels, the foot impulse increased from 14% to 53% of the total load and scaled linearly with strut height adjustments, supporting clinician-prescribed loading increments. Contralateral stance duration increased only modestly compared to commonly used assistive devices, indicating reduced compensatory loading on the intact limb. These findings demonstrate the proof-of-concept feasibility of the OLAFO, highlighting its potential for verifying full offloading and prescribing partial-loading targets during rehabilitation. Future research will evaluate performance across patient populations and clinical rehabilitation tasks.

## 1 Introduction

Various medical conditions affecting the ankle and foot require either complete or partial offloading of the affected region during rehabilitation. These conditions include fractures, sprains, diabetic foot ulcers, Charcot neuroarthropathy, tendon and ligament ruptures, foot deformities, and post-surgical recovery, including ankle fusion or replacement [1–4]. To facilitate ambulation during the recovery period, assistive devices are essential for managing weightbearing and supporting functional mobility [5–8].

Although complete offloading is often prescribed in the early post-injury or post-surgery phase, typically for 4 to 12 weeks [1–3, 9–11], prolonged immobilization and non-weightbearing can lead to secondary complications, including muscle atrophy and bone density reduction in the unloaded leg [12–14]. For certain medical conditions in the later stages of rehabilitation, a gradual transition to partial weightbearing can be beneficial, as it helps preserve bone strength by stimulating osteogenesis, while still meeting clinical requirements for comfort and offloading [3, 9, 10, 15, 16]. In addition, unilateral ankle-foot offloading often leads to gait asymmetries that can reduce gait efficiency, increase energy expenditure, alter kinematics, and increase the incidence of lower back pain [17–21].

Currently, crutches are commonly used to allow patients to walk without loading their affected ankle or foot [22]. However, studies have shown that crutch gait is typically slower and less energetically efficient than normal gait [22–24]. Additionally, crutches limit the functional use of the upper extremities and alter the joint kinematics and ground reaction force (GRF) patterns [24–27]. Importantly, crutch gait increases the loads on the weightbearing leg and upper extremities [24], which can lead to pain, discomfort, and adverse secondary conditions such as stress fractures and hematomas, particularly with long-term use [27–30]. Moreover, crutch users often struggle to adhere to partial weightbearing prescriptions, even after training, which may lead to unintended excessive weightbearing and related complications [31–35].

Alternative devices to crutches include the knee scooter and the hands-free knee crutch, both designed to facilitate distal leg offloading by maintaining the knee flexed at approximately 90°. While both devices can achieve complete non-weightbearing, they differ significantly in their functional performance during gait. The knee scooter is a wheeled aid that provides mobility and stability by positioning the affected limb in knee flexion with the hip extended; however, it requires the use of three of the four extremities. Additionally, prolonged use of the knee scooter may cause adverse secondary effects, including reduced muscle activation and reduced venous flow, which are risk factors for muscle atrophy and deep vein thrombosis [36–39]. In contrast, the hands-free knee crutch enables a more active gait pattern by preserving hip and trunk motion and maintaining cyclic lower-limb muscle activity, while keeping the upper extremities mostly free [24, 40]. Beyond the expected restriction of knee motion, use of this device substantially alters walking kinematics, with compensatory strategies such as increased hip circumduction that can elevate metabolic cost, increase center-of-mass fluctuation, and prolong stance duration of the weightbearing limb [24, 41]. Despite these biomechanical shifts, both the knee scooter and knee crutch are frequently preferred or perceived as less demanding than crutches [24, 39, 42, 43]. However, both devices support the affected limb on an anterior platform at the proximal shank, a configuration that may plausibly contribute to the local discomfort and pain reported with these devices [24, 42, 43].

Another alternative for limb offloading is the use of ankle-foot orthoses (AFOs) and controlled ankle motion (CAM) boots, which redirect GRFs from the plantar surface to the shank, thereby bypassing the injured region. Both AFOs and CAM do not restrict knee and hip motion in the affected leg, which could facilitate greater functional mobility [44, 45]. However, CAM boots and custom AFOs, such as patellar-tendon bearing (PTB) braces and casts, provide only partial and inconsistent offloading and can induce compensatory gait deviations at the knee and hip [44–46].

CAM boots are primarily designed to restrict ankle motion. They can be effective at reducing forefoot pressures, but do not reliably offload the midfoot or hindfoot, and the degree of offloading depends on boot design, sole thickness, and the degree of ankle plantarflexion [44, 45, 47]. Moreover, the added sole height of a CAM boot introduces a leg-length discrepancy, which alters knee and hip kinematics, a mechanism proposed to explain the knee and hip pain reported during and after CAM boot use [44, 48].

PTB-style braces and casts transfer part of the weightbearing load from the distal foot to the proximal tibia and the patellar tendon region, but overall offloading remains partial. PTB braces reduce overall plantar pressure by approximately 20–48%, and can paradoxically increase focal forefoot pressure [46, 49]. The Böhler-Walker frame (Beagle Orthopaedic, Blackburn, UK) reduces metatarsal head forces by approximately 60–70%, substantially outperforming the Sarmiento PTB cast [50]. PTB-type devices suffer from limited comfort, require specialized fabrication and expert fitting, and produce atypical gait patterns that reduce patient acceptance [46, 50, 51]. Furthermore, most AFOs and casts elevate the foot to prevent plantar contact, requiring contralateral shoe elevation to restore leg-length symmetry [46, 48].

For diabetic foot ulcers, a variety of therapeutic footwear and insole designs have been developed to offload high-risk plantar regions by redistributing pressure to adjacent plantar areas, either passively or actively [5, 6, 52, 53]. However, even with optimized designs, these solutions often fail to provide sufficient pressure reduction at all high-risk sites. This offloading often shifts the load to neighboring plantar regions, which may also experience elevated pressures, thereby increasing the risk of injury in those new locations [5, 6].

Most ankle-foot orthoses (AFOs) are custom-designed to individual anatomy and fitted in specialized clinics. However, the ZeroG AFO (Certified Orthopedics, Inc., Fort Collins, CO, USA) is marketed as the only prefabricated brace that provides complete offloading [54]. In a recent comparative gait study, the ZeroG AFO showed favorable biomechanical and metabolic energy performance compared to other complete-offloading devices, but important limitations were also identified, including poor comfort rating, excessive pressure and pain around the shank region, atypical ankle kinematics, and difficulty in preventing forefoot contact during late stance [24].

Given these limitations of existing assistive devices, this study presents the design and evaluation of a novel patent-pending offloading AFO (OLAFO) [55]. The OLAFO is designed to facilitate both complete offloading and adjustable levels of partial offloading. It incorporates a patient-specific load-bearing shank brace that distributes load across the shank surface, and mediolateral side struts that facilitate adjustable load sharing between the shank and the foot. Additionally, mediolateral contact plates are designed with a roll-over rocker profile, derived from gait measurements, to promote a more physiological walking pattern. This study describes the OLAFO design and fabrication methodology and evaluates its performance in an able-bodied participant during complete offloading and five partial loading levels. The evaluation quantifies the GRF and impulses transmitted to the affected foot at each loading level, and the stance phase (SP) symmetry index (SI).

## 2 Methods

### 2.1 OLAFO Design Methodology

The OLAFO consists of three primary components:

1. A patient-specific load-bearing shank brace (section 2.1.1);
2. Two adjustable side struts (section 2.1.2);
3. Two mediolateral contact plates (section 2.1.3).

Figure 1 illustrates these components, both in an assembled view and an exploded view, which shows the leg positioning within the device.

**Figure 1.**
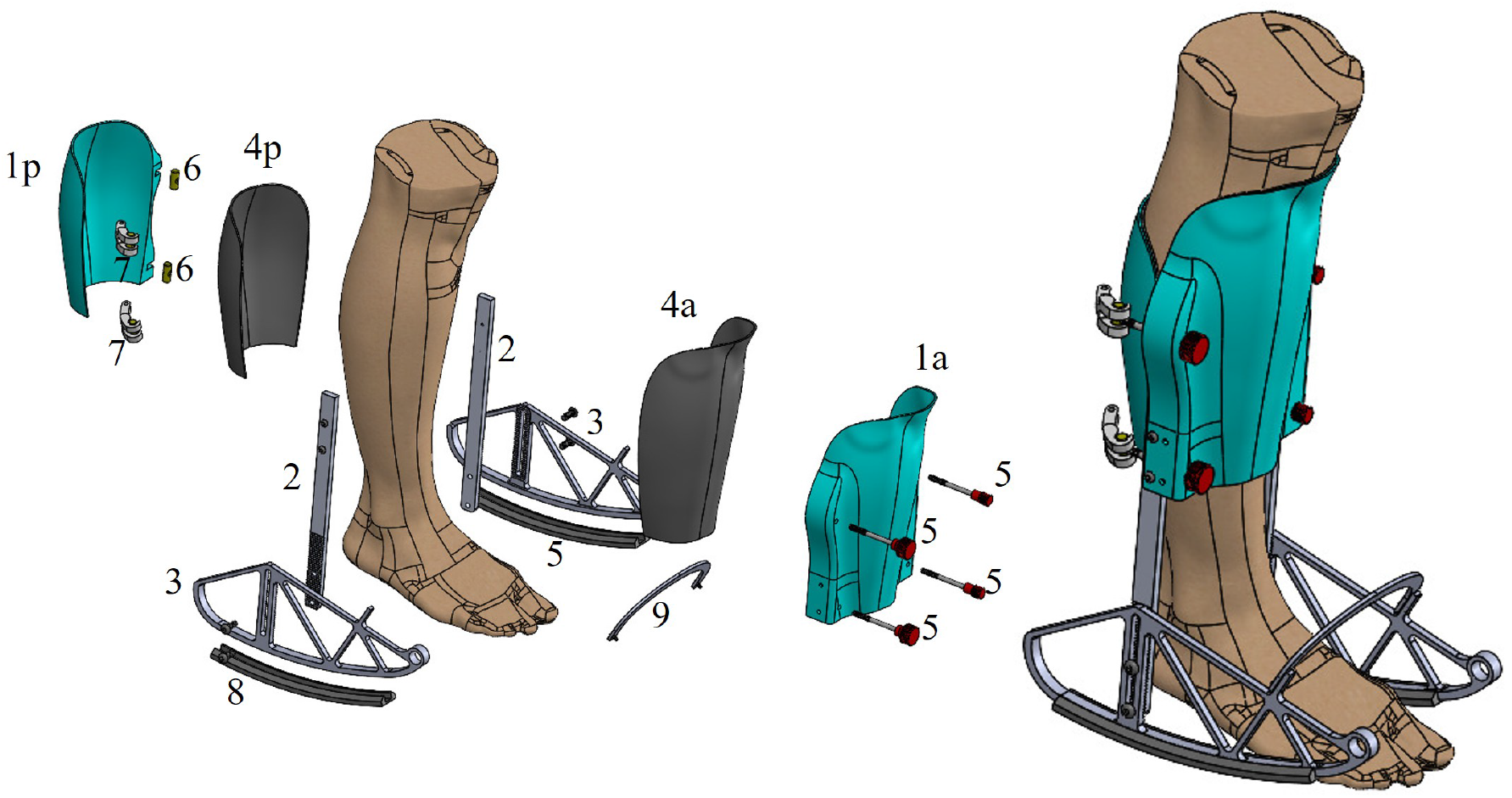
An assembled (right) and exploded (left) view of the OLAFO showing its components: 1a - anterior shank brace; 1p - posterior shank brace; 2 - adjustable side struts; 3 - mediolateral contact plates; 4a/4p - anterior and posterior 5 mm neoprene liner (SEPA DIVER, Trieste, Italy); 5 - screws with knurled knobs; 6 - barrel nuts; 7 - catch hooks; 8 - TPU outsole tread; and 9 - dorsal bridge.

#### 2.1.1 Patient-Specific Load-Bearing Shank Brace

The custom load-bearing shank brace is a crucial component of the OLAFO design, which serves as the mechanical interface with the patient’s soft tissues. Its primary function is to distribute the load across the shank to maintain comfort and tissue integrity.

To construct a digital model of the participant’s lower leg, we used an EinScan-H handheld 3D scanner (Shining3D Technology, Hangzhou, China), which has been reported to achieve sub-millimeter accuracy [56, 57]. The scanning was performed with the knee in a natural standing posture and the foot offloaded by shifting the entire weight to the contralateral leg.

The scanning process captured both the surface geometry and color texture of the patient’s limb, enabling us to mark the anatomical regions required for brace modeling (Figure 2a). The landmarks for the knee and ankle joint axes were used to align the model within the anatomical planes. The bony regions of the fibula head and anterior tibia, as well as the patellar tendon region, were marked to guide the brace rectification and cutlines, using a methodology inspired by the design of transtibial prosthetic sockets [58–60]. The scanning process resulted in a triangular mesh with texture that represents the outer surface of the lower leg, which was exported in OBJ format (Figure 2a).

**Figure 2.**
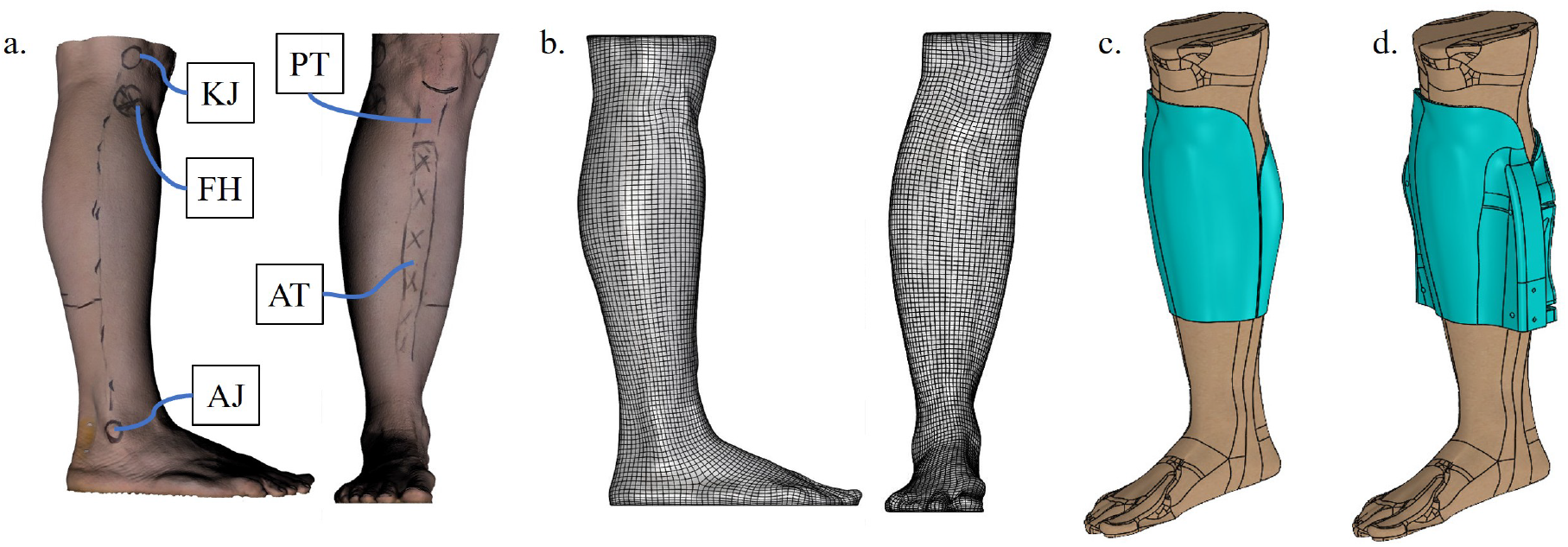
Main steps of shank brace design. (a) Lateral and anterior views of the 3D scan mesh, showing marked anatomical regions of interest: knee joint (KJ), ankle joint (AJ), fibula head (FH), anterior tibia (AT), and patellar tendon (PT). (b) Parametric subdivision surfaces generated from the triangular mesh. (c) CAD brace model following the generation of cutlines and rectifications. (d) Final CAD after modeling the side connectors and fastening interfaces.

To initiate the computer-aided design (CAD) workflow, the mesh was converted into a parametric subdivision (SubD) surface using Rhinoceros 7 (Robert McNeel and Associates, Seattle, WA, USA) (Figure 2b). Based on the drawn anatomical landmarks, the fibula head and anterior tibia regions were rectified by extruding outwards by 3 mm, followed by tangential blending with the adjacent surfaces. The rectified leg model was exported in STP format and imported into SolidWorks 2025 (Dassault Systèmes, Vélizy-Villacoublay, France), where the remainder of the design process was carried out.

The proximal brace cutline was defined using the anatomical landmarks drawn on the patellar tendon, posterior shank, and medial-lateral knee joint, which served as control points for a cubic spline representation [58, 61]. The patellar tendon was intentionally included beneath the brace, as it is considered pressure-tolerant and is conventionally used as a primary load-bearing region in PTB braces and prosthetic sockets [62–65]. In contrast, the regions over the knee joint and proximal posterior calf were excluded from the brace to permit comfortable knee flexion. The distal cutline was positioned 180 mm above the floor level (Figure 2c). Alternative heights may be chosen according to injury location or clinical requirements, for example, to accommodate a cast, external fixation, or compression bandaging.

The brace model was then divided into anterior and posterior shells to facilitate donning and tightening. The separation line between the two shells runs along a line connecting the ankle and knee joints on both the medial and lateral sides (Figure 2c). Next, a 2 mm outward offset was applied to both shells to account for the thickness of the compressed Neoprene liner between the skin and the brace. Finally, the shells were given a 2 mm thickness to obtain a volumetric model, and the medial and lateral side connectors were integrated to allow interfacing with the adjustable side struts (Figure 2d).

The brace donning and tightening mechanism was designed to enable fast, secure, and adjustable fastening. Two adjustable hinges and buckles are positioned on the medial and lateral sides of the brace, respectively, as shown in Figure 1 (components 5-7). The buckles allow opening and closing of the brace around the leg, while the hinges prevent vertical translation of the posterior shell. Once the brace is donned, its tightness can be independently adjusted at each of the four knobs without reopening the buckles, allowing the brace to be doffed and re-donned while maintaining the same tightness level.

Next, the brace was manufactured using a hybrid method that combines a 3D-printed core with external carbon fiber composite lamination. The core was 3D-printed from polylactic acid (PLA) filament (PLA+, eSUN Industrial Co., Ltd., Shenzhen, China), as shown in Figure 3a. To increase strength and stiffness, the core was externally reinforced with three layers of 208 g*/*m^2^ carbon fiber sheets oriented at 45^*°*^ relative to the tibia axis. The carbon fiber layers were bonded using a Biresin CR80 epoxy resin combined with a CH 80-2 hardener (Sika Services AG, Zurich, Switzerland).

**Figure 3.**
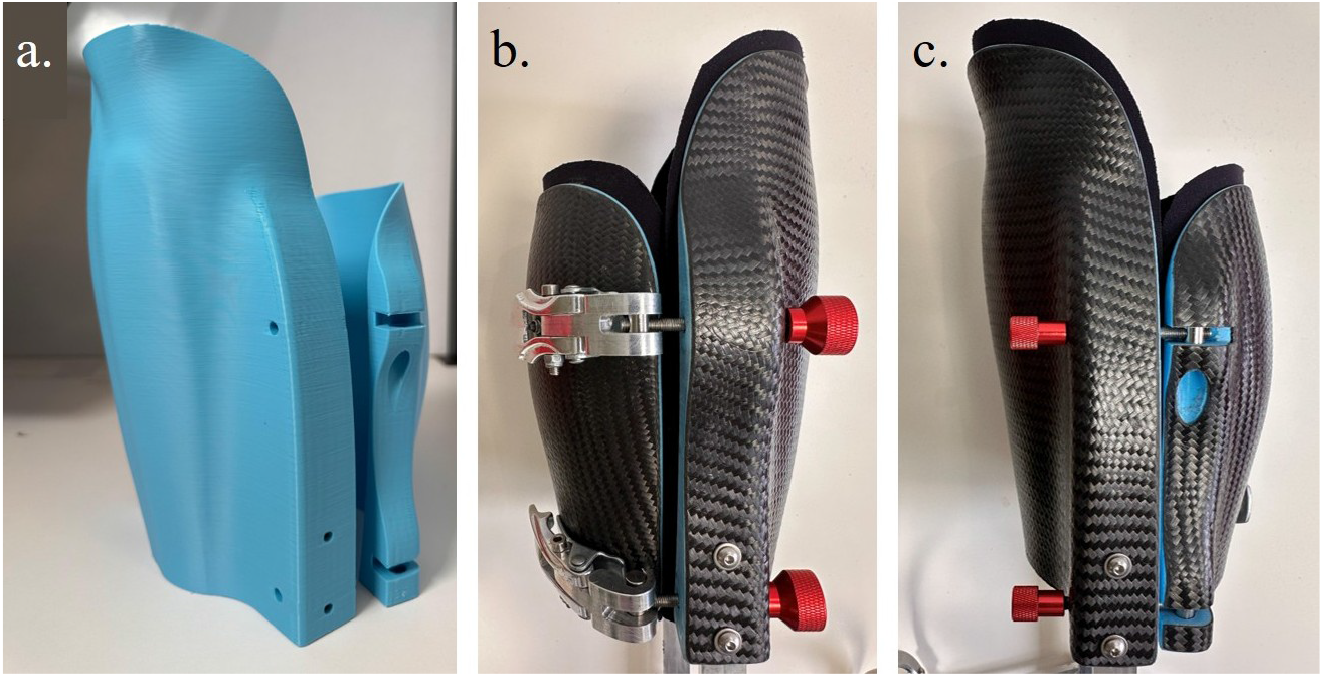
OLAFO brace fabrication. (a) 3D-printed PLA core of the brace. (b) Lateral view of the brace after carbon-fiber composite lamination, showing the buckles used for donning and tightening. (c) Medial view showing the integrated adjustable hinges.

A 5 mm thick Neoprene rubber sheet (SEPA DIVER, Trieste, Italy) was glued to the internal brace surface as a liner, providing cushioning and absorbing shear deformation between the brace and the skin. This method was preferred over a sleeve liner, which may be difficult to don over an injured foot or ankle. Finally, two buckles and two hinges, each featuring a screw with a knurled knob, were assembled on the medial and lateral sides of the brace, respectively (Figure 3b-c).

#### 2.1.2 Adjustable Side Struts

The amount of load transferred to the injured foot is adjusted by changing the height of the plantar surface above the ground. This height adjustment is achieved using interlocking serrated struts, which can be fastened together in discrete increments of 2 mm, up to a maximum of 50 mm, as shown in Figure 4a. These struts are connected proximally to the shank brace and distally to the contact plates (Figure 4b-c). In a specific configuration, the strut length determines the vertical offset between the bottom surface of the contact plates and the plantar surface. When the foot is not in contact with the ground, 100% of the load is transmitted to the shank through the side struts and the brace (Figure 4b). However, when the plantar surface is in contact with the ground (Figure 4c), tissue deformation allows the load to be shared between the shank and plantar surfaces at different ratios, depending on the adjusted height.

**Figure 4.**
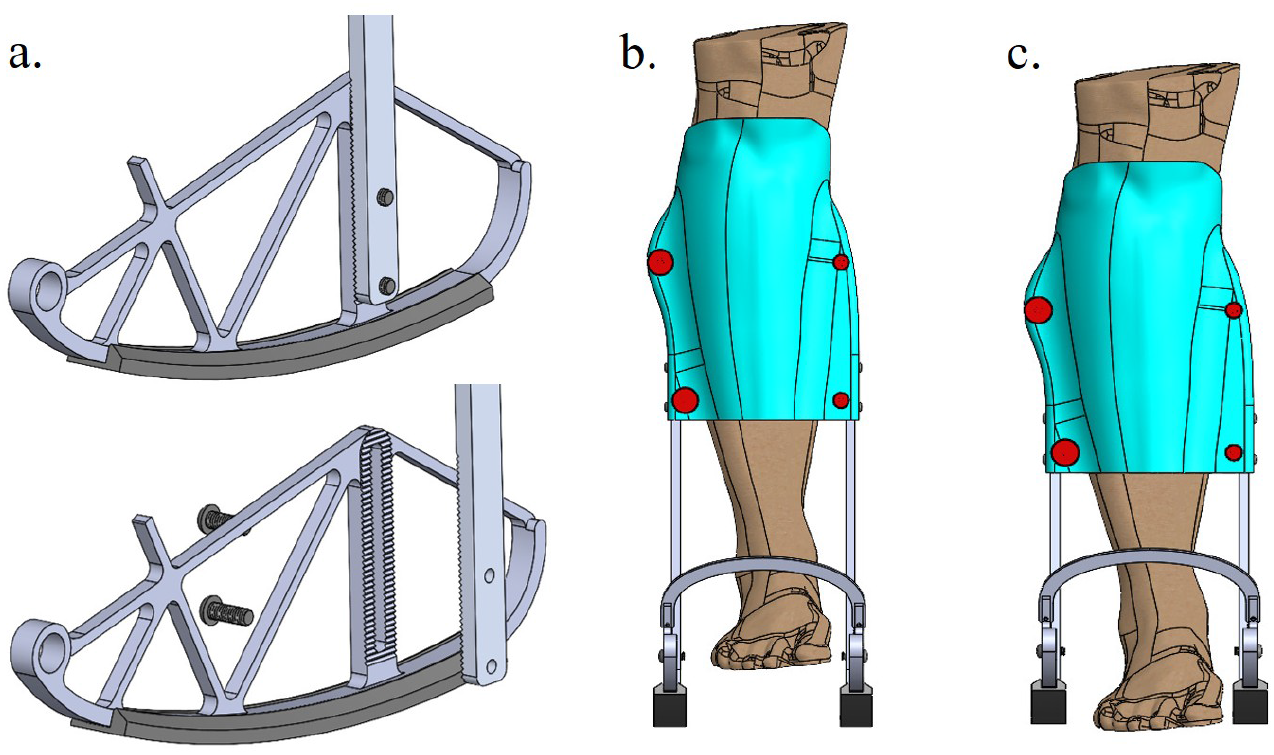
Adjustable side struts mechanism and loading configuration. (a) Zoomed-in view of the adjustable interlocking serrated struts and screw-locking, highlighting the fine-tuning capability. (b) Complete offloading configuration, in which the plantar surface is elevated to the maximum height and is not in contact with the ground, thus completely offloaded. (c) Partial loading configuration, in which the plantar surface is in contact with the ground surface, allowing distribution of the GRF between the plantar surface and the shank surface.

#### 2.1.3 Mediolateral Contact Plates

The OLAFO includes two contact plates positioned medially and laterally with respect to the foot. Unlike designs in which the sole is beneath the foot, this design allows the foot to remain close to the ground, thereby reducing the need to elevate the contralateral foot to maintain symmetry. In the complete offloading configuration, the affected foot can be positioned at the same distance from the ground as the shoe sole thickness of the contralateral leg. To facilitate a natural and symmetric walking pattern, we used rocker soles to compensate for the device’s lack of ankle flexion. The rocker profiles were designed according to the ankle-foot roll-over-shape (ROS), which describes the effective geometry to which the ankle-foot conforms from initial contact (IC) to opposite initial contact (OIC) [66]. The ankle-foot ROS can be quantified by computing the foot center of pressure (CoP) trajectories in a coordinate system fixed to the shank [66]. Typically, these trajectories are then fitted with a circular arc [66–68].

Figure 5 illustrates the transformation of the knee and ankle marker trajectories as well as the CoP in the sagittal plane during one SP from the laboratory’s global coordinate system (CS) {x,y,z} (autoreffig: ROSa) to a local CS {u,v,w} fixed to the shank, with the origin at the ankle joint (Figure 5b). The transformed CoP trajectories from IC to OIC were fitted with a circular arc using the methodology described by Hansen et al. [66, 67]. After obtaining the ROS arc, it was integrated into the design of the contact plates.

**Figure 5.**
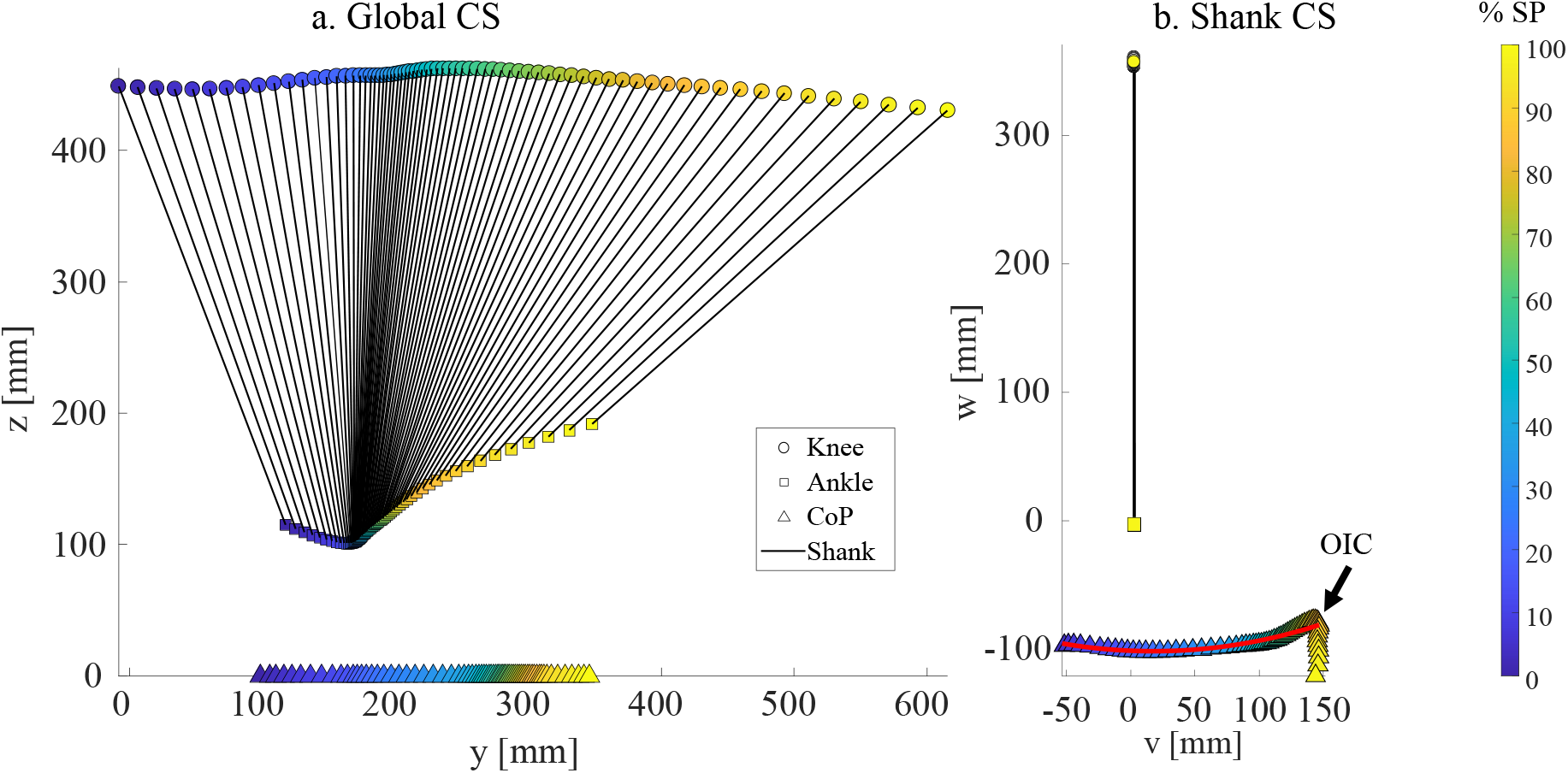
Computation of Roll-over-shape (ROS) from walking measurements. (a) Sagittal plane trajectories of lateral knee and ankle markers, and the foot center of pressure (CoP) in the global laboratory coordinate system (CS) (y-z) during the stance phase (SP). (b) Trajectories transformed into the shank-fixed CS (v-w) with the origin at the ankle joint. The marker colors indicate the percentage of SP. The red curve represents the circular arc (radius of 430 mm), fitted to the transformed CoP from initial contact (IC) (0%SP) to opposite initial contact (OIC), which occurred at 88%SP.

A 3D-printed outsole made of thermoplastic polyurethane (TPU) was designed to cover the distal part of the contact plates, and a rubber tread was glued to the bottom surface to enhance grip and cushioning (Figure 6a). Additionally, a dorsal bridge was incorporated between the medial and lateral plates to increase the structure’s stiffness. This bridge also provides a mounting surface for straps that secure the foot in place and prevent unwanted plantarflexion that could lead to foot-ground contact (Figure 6b-c). The contact plates and dorsal bridge were fabricated using CNC machining from aluminum alloy 6061, resulting in a combined mass of 225 g. Together with the brace and fasteners, the OLAFO’s total mass was 1170 g.

**Figure 6.**
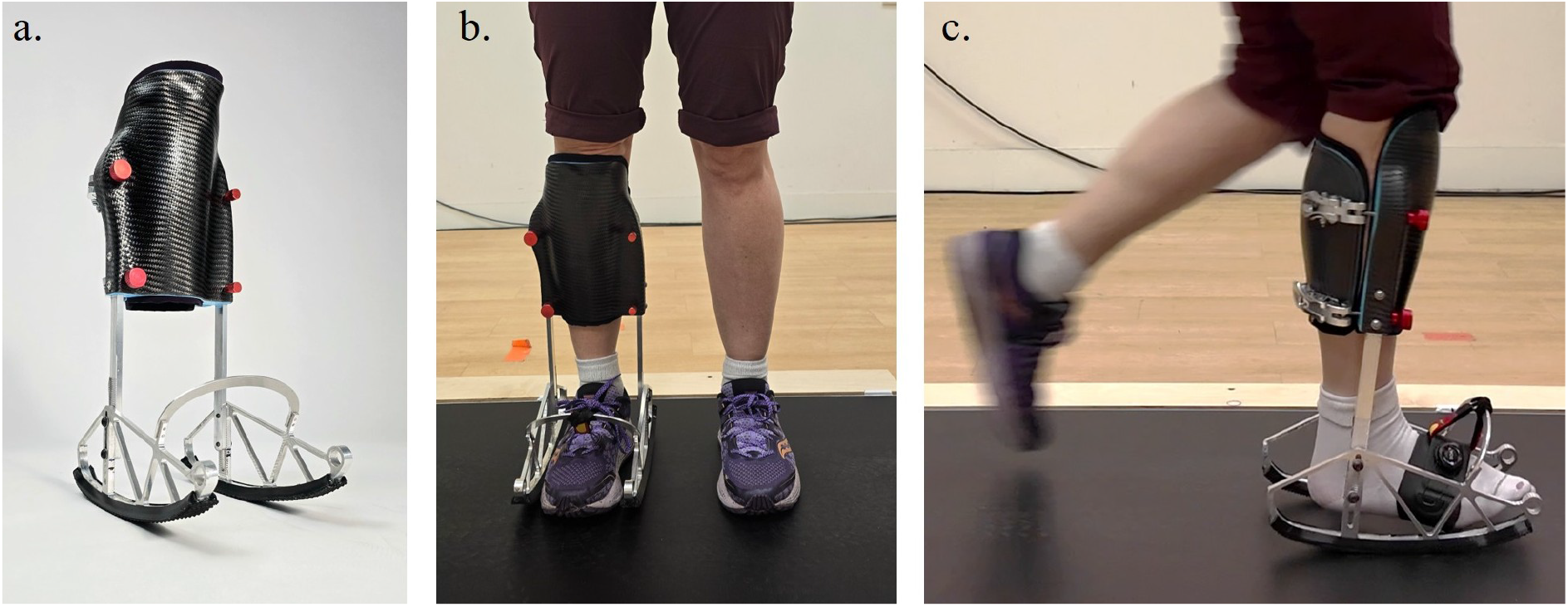
(a) Perspective view of the fully assembled OLAFO device. (b) The OLAFO donned on the participant’s right leg. (c) Walking on the pressure-instrumented walkway with the OLAFO in a complete offloading configuration, with a strap helping suspend the foot from the OLAFO’s dorsal bridge.

### 2.2 Human Subject Evaluation

#### 2.2.1 Experimental Protocol

An able-bodied participant was recruited for this study (female, age: 41, height: 160 cm, mass: 62 kg). The study was approved by the Technion Institutional Review Board (#108-2020) and the participant provided written informed consent prior to participation. The protocol involved two visits. In the first visit, the participant’s right leg was 3D-scanned to inform the design of the custom brace, as described in section 2.1.1. Additionally, the participant completed 10 walking trials at self-selected speed along a 10 m straight walkway. A 16-camera motion capture system (Vicon Motion Systems Ltd, Oxford, UK) was used to track lateral knee and ankle joint markers at 120 Hz, and two floor-embedded force plates (OR6-7-1000, AMTI Inc., Watertown, MA, USA) recorded GRF and CoP data at 960 Hz. These data were used to inform the contact plate’s ROS, as described in section 2.1.3.

Following device fabrication and assembly (fig: liveAFOa), the second visit took place. Dur-ing this visit, the OLAFO was fitted to the participant’s right leg, as shown in Figure 6b. After familiarization with the device, gait evaluation was conducted under 7 conditions: complete offloading, 5 levels of partial loading (side struts adjusted in 4 mm increments), and a control condition involving normal walking without the device. All conditions were conducted at the participant’s self-selected speed.

A pressure-instrumented walkway (FDM-2, Zebris Medical GmbH, Isny im Allgäu, Germany) was used to record plantar pressure data at 200 Hz. Between 10 and 15 gait cycles (GCs) were recorded for each condition. During the normal gait control condition and partial loading conditions, the participant walked with both shoes on. In the complete offloading condition, the right foot (OLAFO side) was tested without a shoe, and the height of the side struts was set to match the shoe sole thickness, so that the plantar surface was at approximately the same height from the floor on both sides (see Figure 6c). For complete offloading, a strap suspended the foot from the dorsal bridge (see Figure 6c), whereas during partial loading conditions, the strap was connected to the shoelaces.

#### 2.2.2 Data Analysis

The recorded pressure data were extracted from the individual sensor readings using the Zebris walkway software (Zebris FDM 3.0.14) and processed in MATLAB (R2022b, The MathWorks, Inc., Natick, MA, USA). For the complete offloading trials, the OLAFO side was segmented into two regions: medial and lateral contact plates (Figure 7a), whereas for the partial loading levels, it was segmented into three regions: medial and lateral contact plates, and the foot between them (Figure 7b). The intact foot was defined as a single region. For each region, the vertical GRF was computed by summing the products of pressure and sensor area across the corresponding region:

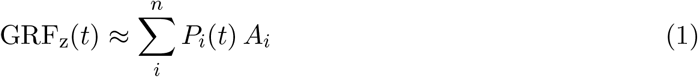

where GRF_z_ is the vertical GRF at time *t, P*_*i*_(*t*) is the pressure measured by the *i*^*th*^ sensor at time *t, A*_*i*_ is the area of the *i*^*th*^ sensor, and *n* is the total number of sensors in the region.

**Figure 7.**
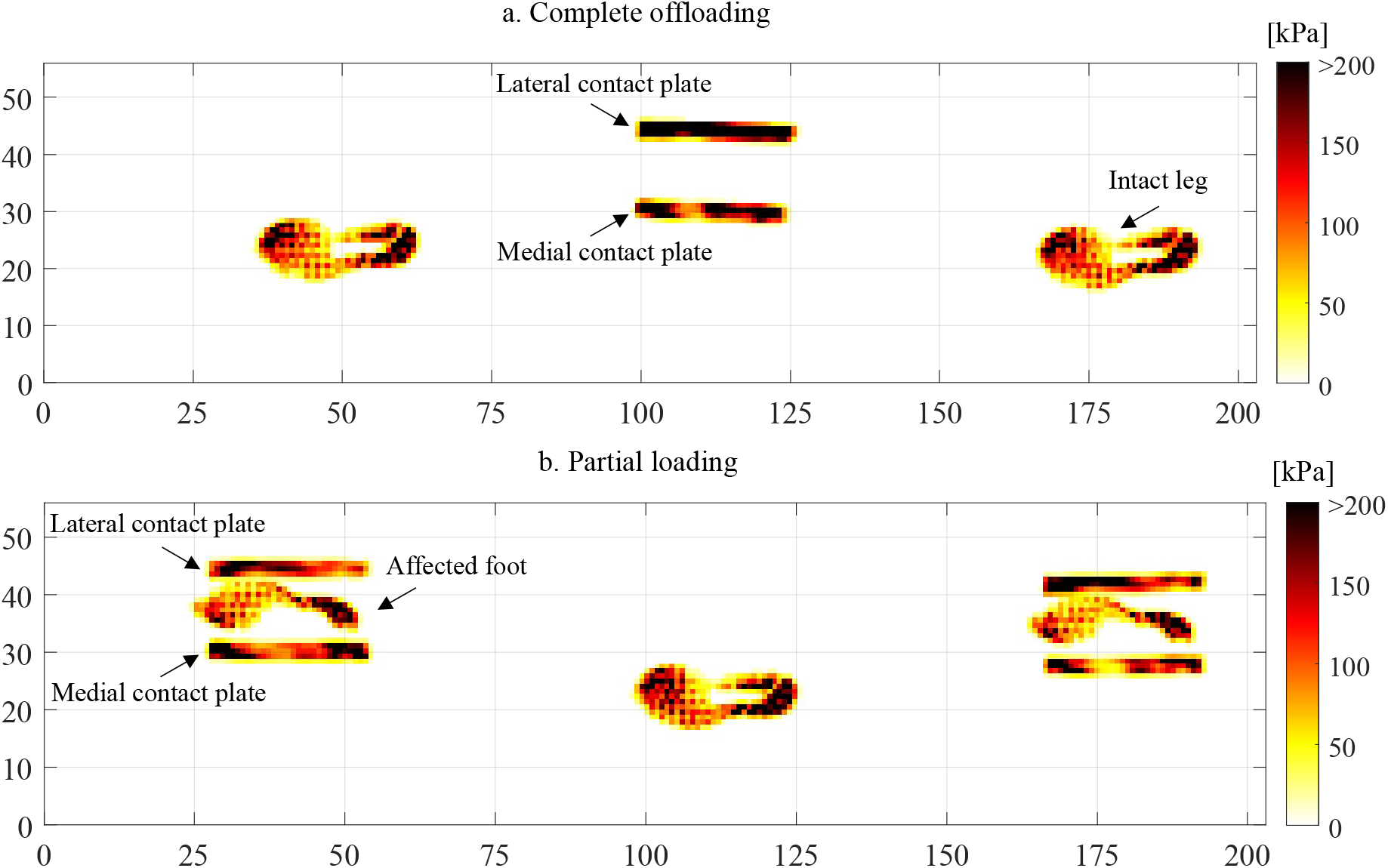
Representative maximum pressure maps recorded during walking with the OLAFO: (a) complete offloading condition, showing pressures under the contact plates and the intact foot; (b) partial loading condition, showing pressures under the contact plates, the affected foot, and the intact foot. The axes represent the walkway dimensions in centimeters.

Gait events, including IC and toe-off (TO), were identified from the vertical GRF profiles using a threshold of 1% body weight (BW). GCs in which the participant stepped on the walkway’s edges were excluded from the analysis. For each valid trial, the SP was defined from IC to TO for either the OLAFO or the intact side. All data were temporally aligned to 100% of SP using linear interpolation.

The impulse of GRF_z_ was computed for each region using trapezoidal integration over time. The ratio between the impulse exerted on the foot and the impulse exerted on the entire leg (foot + OLAFO) was computed to provide an overall measure of loading ratio for the partial loading conditions.

Furthermore, the SI of the SP was calculated using:

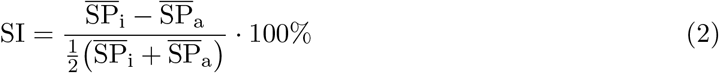

where 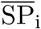 and 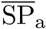 are the mean SP durations of the intact (left) and the affected (right) legs, respectively.

## 3 Results

Following the first visit, analysis of the participant’s walking trials resulted in an ROS with a fitted circular arc of radius 430 mm, as shown in red in Figure 5b. This arc was incorporated into the design of the OLAFO’s contact plates. Once the OLAFO was fabricated and assembled, the participant could easily don and doff the shank brace and adjust its tightness. Additionally, the participant successfully maintained a single-leg standing with the OLAFO, with the foot completely non-weightbearing with no visible limb slippage of the shank within the brace. During the acclimation period, the participant was able to walk with complete offloading and reported a natural and comfortable walking experience.

The evaluation results from the walking experiments are presented below. We first describe pressure distribution maps during walking with complete and partial offloading, then present vertical GRF waveforms and foot loading impulses across strut heights, and SP SI.

fig: MPPa and b present representative maximum pressure values measured during complete and partial offloading, respectively. Figure 7a indicates that the foot generated no contact pressures, and the entire GRF was transmitted through the contact plates. In contrast, Figure 7b illustrates how the GRF is shared between the foot and the device, resulting in partial loading of the foot.

Figure 8 presents the vertical GRFs waveforms for the intact leg, OLAFO leg, medial and lateral contact plates, affected foot, and normal gait control. During complete offloading (Figure 8a), the total OLAFO GRF exhibited a lower and temporally delayed first peak compared with normal gait. As side-strut height decreased from partial loading 1 to 5, the affected foot GRF monotonically increased while the contact plates’ GRFs decreased. In addition, the first peak of the total OLAFO GRF gradually shifted towards the magnitude and timing observed in normal gait (Figure 8b-f). Concurrently, the affected foot GRF evolved from a flat pattern to a characteristic two-peak pattern that increasingly resembled normal gait. The intact leg’s vertical GRF preserved its characteristic pattern. The first peak increased in all conditions by an average of 12%BW above the normal gait value, whereas the second peak remained comparable to normal gait.

**Figure 8.**
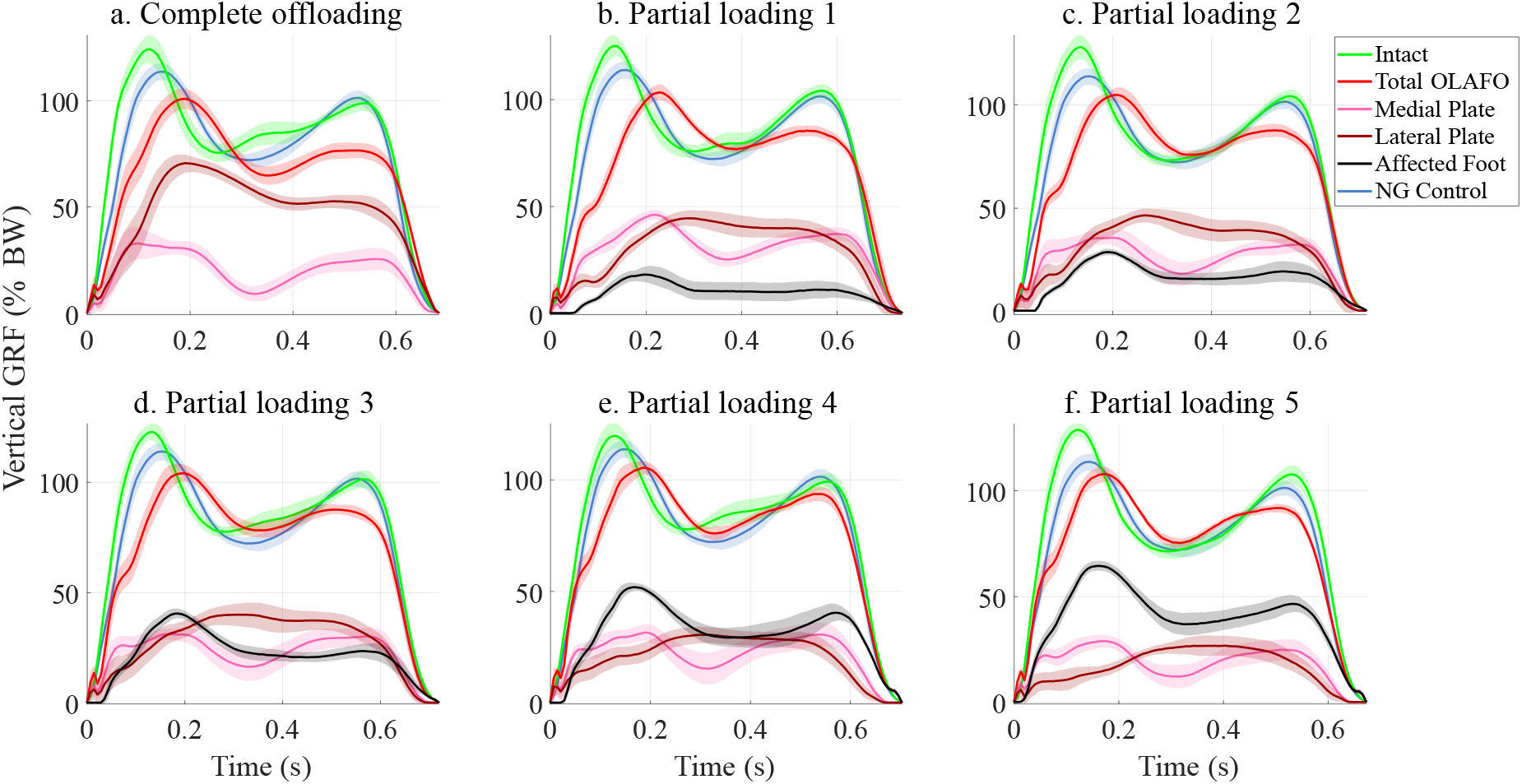
Vertical ground reaction force (GRF) waveforms, normalized by body weight (BW), during walking at self-selected speed in complete and partial offloading conditions. (a) Complete offloading condition, showing vertical GRFs under the intact foot, the total OLAFO (resultant sum of contact plates), and the medial and lateral contact plates. (b-f) Partial loading conditions, with five levels of side-strut height, from highest (partial loading 1) to lowest (partial loading 5). The plots show the vertical GRFs under the intact foot, total OLAFO (sum of contact plates and affected foot), medial and lateral contact plates, and the affected foot. For all conditions, the vertical GRF of unassisted normal gait (NG) control is overlaid in blue for comparison. Solid lines denote median waveforms, and shaded regions show the range across all recorded steps.

Figure 9a shows the vertical GRF on the affected foot as a function of SP percentage for all side-strut height levels, enabling direct comparison across conditions. In the complete offloading condition, the affected foot carried no load. As the strut height decreased from partial loading 1 to 5, loading on the affected foot progressively increased throughout the SP.

**Figure 9.**
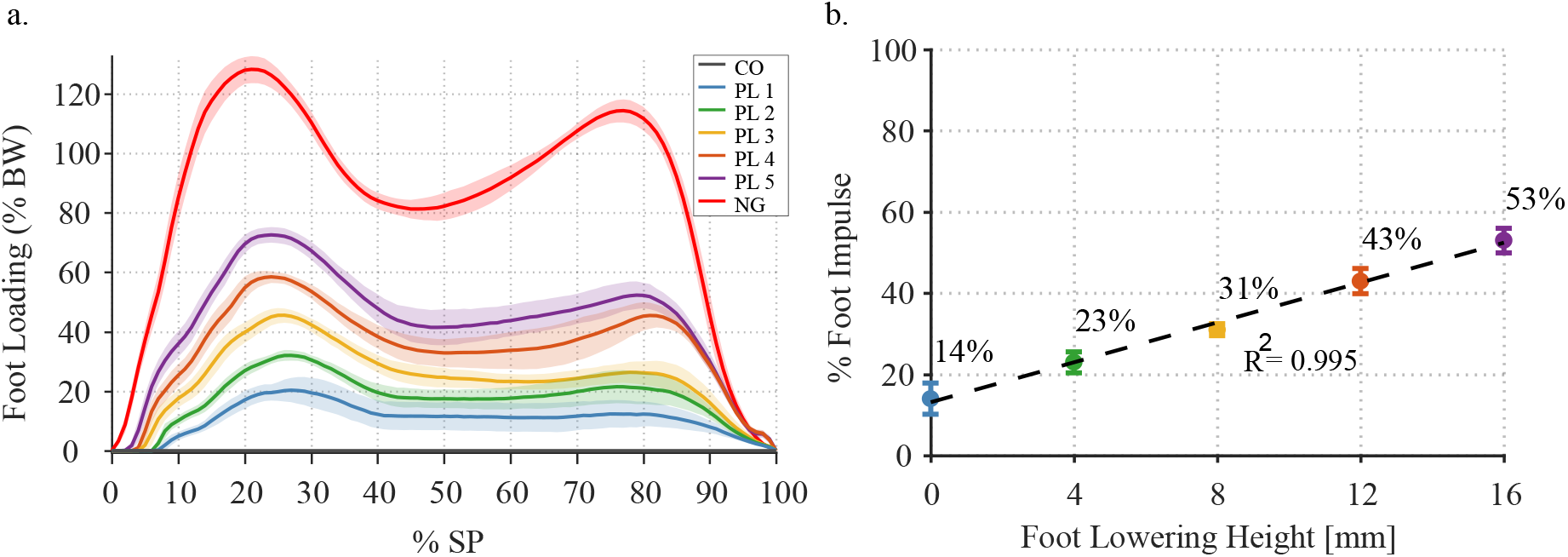
Foot loading with the OLAFO across conditions. (a) Vertical GRF waveforms of the affected foot, normalized by body weight (BW), across all conditions as a function of stance phase (SP): complete offloading (CO), five partial-loading (PL) levels, and normal gait (NG) as a baseline. Solid lines indicate the median waveforms, and shaded regions indicate the range across all steps. (b) Foot loading impulses across the experimental conditions: CO and PL1-PL5, corresponding to foot lowering levels of 4 to 16 mm. The plot shows the mean foot loading impulse as a percentage of the total OLAFO load, with error bars representing the standard deviation across trials. The dashed line represents a linear regression fitted to the PL data (*R*^2^ = 0.995).

Figure 9b presents the affected foot impulse as a percentage of the total impulse on the affected leg for the five partial loading levels. As the foot was lowered in four increments of 4 mm each, this percentage monotonically increased from 14% to 53%. A linear regression fitted to these data yielded *R*^2^ = 0.995.

Figure 10 presents the SP durations and resulting SI values for the intact and OLAFO-assisted legs across all conditions. In the complete offloading condition, the OLAFO leg exhibited a shorter SP (58.1%GC) than the intact limb (65.9%GC), yielding the largest SI (12.57%). In the partial loading conditions, the asymmetry decreased compared to complete offloading. The greatest SI (10.83%) was observed at the highest strut-height level (i.e., greatest offloading), and smaller values between 6.15-8.38% were obtained for the other partial-loading levels, alongside OLAFO-side SP durations between 59.2-60.2%GC. In the normal gait condition, SP durations were nearly identical for both limbs (63.7 vs. 63.1%GC), yielding an SI of -0.96%.

**Figure 10.**
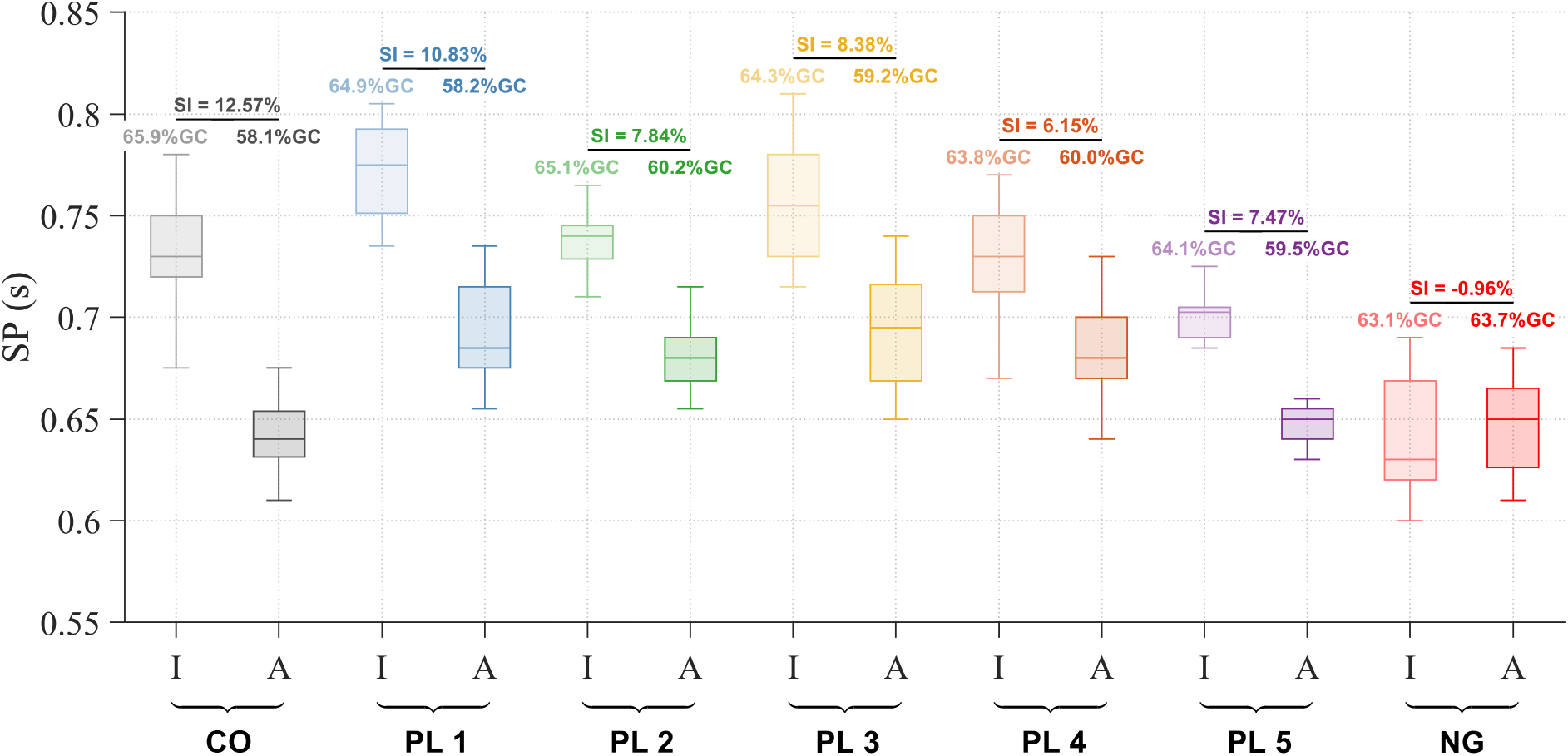
Stance phase (SP) durations for each limb (Intact (I) and OLAFO-assisted (A)) across all conditions: complete offloading (CO), five partial-loading (PL) levels, and normal gait (NG). The percent of gait cycle (GC) corresponding to each limb and condition is indicated above the boxes. Each box shows the median, quartiles, and range of SP durations across all trials. For each condition, the symmetry index (SI) (Eq. (2)) of the SPs is shown above the corresponding horizontal line.

## 4 Discussion

This study presented an innovative OLAFO, comprising three main components: a patient-specific load-bearing shank brace, adjustable side struts, and mediolateral contact plates. A preliminary proof-of-concept evaluation was conducted with an able-bodied participant walking with the OLAFO, quantifying vertical GRFs under complete offloading and five partial-loading levels, the corresponding affected foot impulses, and the SI of the SP.

The shank brace was developed using a digital workflow, providing a repeatable, data-driven alternative to traditional AFO fabrication. Unlike conventional methods that rely on expert casting and time-consuming manual molding, this approach improves consistency and repeatability and can be more easily standardized across clinics [69–71].

The customized load-bearing shank brace, along with its fastening system that enables quick donning and adjustment of tightness, has shown practical benefits in ease of use and comfort. In our evaluation, the participant was able to don and tighten the brace quickly, without reports of shank discomfort, and to doff and re-don it at the same tightness, without loosening the buckles. These features address previously reported challenges with offloading AFOs that rely on prefabricated braces and strap-based tightening, which can be time-consuming to don and adjust [24], can cause local discomfort and pain around the shank [24], and PTB braces that can be sensitive to improper or loose donning [51].

The side struts enabled adjustability of the load-sharing ratio between the foot and the shank. As the strut height decreased, the affected foot exhibited progressively higher loading and a GRF pattern more typical of non-disabled gait, including restoration of clear initial loading and push-off peaks (Figure 8 and Figure 9). This progression suggests that the participant is experiencing increasing physiological loading on the affected side, which is desirable during graded rehabilitation. Such gradual loading is clinically important, as appropriate mechanical stimulation has been shown to help maintain bone strength and support soft-tissue healing [16, 72]. In contrast to the OLAFO, patients who use crutches often struggle to maintain prescribed partial weightbearing consistently during gait [32–35]. In this study, we focused on offloading levels where the foot load is reduced by at least 50%BW, because this value lies at the upper end of clinically used partial-weightbearing ranges, whereas smaller reductions leave plantar forces relatively close to full loading and are unlikely to justify the added mass and bulk of a knee-high brace [6, 73].

The ROS-informed rocker profile of the contact plates was derived from the participant’s able-bodied gait and is therefore patient-specific in this proof-of-concept. For patients whose medical condition precludes reliable measurement of a ROS, parameterized models from the literature could be used [66, 67, 74]. These models provide generalized ROS parameters that approximate typical gait dynamics and can serve as a basis for functional contact plate geometries when patient-specific data are unavailable. Alternatively, predictive musculoskeletal simulations could be used to determine the optimal rocker design [75].

Overall, the OLAFO supported a natural walking experience under both complete and partial offloading conditions. In the complete offloading condition, the maximum pressure maps (Figure 7) show no plantar contact under the affected foot, confirming correct donning and full weightbearing through the shank brace. This addresses limitations reported for other AFOs and braces. For example, with the prefabricated ZeroG AFO and the custom Toad Anti-Gravity Brace (Toad Medical Corporation, Carson City, NV), avoiding forefoot contact in late stance was challenging [24, 76]. Several other AFOs, braces, and casts were unable to achieve complete offloading, with maximum offloading typically in the 30% - 90% range [5, 6, 45, 50, 76, 77]. Additionally, issues of leg slippage within the brace have been reported for PTB braces [51].

The slight elevation of the first peak of the contralateral leg vertical GRF compared to the control condition (Figure 8) indicates that the intact limb accepts slightly more load during weight acceptance, reflecting a compensation strategy rather than perfectly symmetric loading. However, because this elevation remained modest and its timing closely matched the normal gait baseline, the OLAFO appears to avoid the pronounced overcompensation often observed with other devices [24, 27, 78, 79].

The differences between the OLAFO GRFs and those of the intact limb and normal gait likely reflect both altered ankle mechanics and load transfer within the device. While the ROS profile was designed to reproduce the CoP trajectories in the shank CS, it does not replicate the normal storage and rapid release of elastic energy in the Achilles tendon and intrinsic foot structures, so push-off is generated more by a smooth rolling motion than by elastic recoil, which lowers and broadens the second GRF peak [80–82]. In addition, walking with a new assistive device may lead to more cautious foot placement and promote an impact-reduction strategy; such strategies have been shown to lower the first vertical GRF peak and loading rate without necessarily changing walking speed [83]. Finally, because a substantial portion of the limb load is transmitted proximally through the shank brace, with deformation of shank soft tissues and brace liner, some impact is absorbed away from the plantar region, which can further smooth and reduce the resultant GRF peaks.

The linear relationship observed between side-strut height and the relative foot impulse (Figure 9) provides a simple and clinically useful control parameter. This mapping enables clinicians to select target partial-loading percentages and prescribe precise weightbearing incre-ments. Such control can help structure the progression from complete offloading to gradually increasing partial loading during rehabilitation, a process reported to be difficult to achieve consistently with other devices [32–35], with many patients exceeding prescribed limits even with training.

In the complete offloading condition, the contralateral SP duration increased to 65.9%GC, compared to 63.1%GC during normal gait (Figure 10). This increase is smaller than previously reported for other devices, including the ZeroG AFO (from 62 to 68%GC), the knee-crutch (from 62 to 72%GC), and forearm crutches (from 62 to 76%GC) [24]. This finding suggests that the OLAFO may provide better stability on the affected leg and require less compensatory loading on the intact leg than other assistive devices, although this observation is based on a single participant and cannot be evaluated statistically.

The SP durations and corresponding SI results (Figure 10) show that SP timings are more symmetric at partial loading than in complete offloading. Larger SI in complete offloading indicates that the intact leg prolongs stance duration when the affected foot is fully unloaded, consistent with a compensatory strategy reported for unilateral push-off impairment [18, 84]. As more load is transferred to the affected foot, SP on the OLAFO leg lengthens toward intact-limb and normal gait values, and SI values move closer to the near-zero value observed in normal gait, indicating a partial restoration of temporal symmetry. Notably, even under complete offloading, the SI remained below 13%, indicating only a modest degree of SP asymmetry.

This study has several limitations. First, this proof-of-concept evaluation was conducted in a single able-bodied participant, which precludes any statistical inference and limits the generalizability of the findings to broader patient populations. Second, ROS is often difficult to reliably measure in patients with ankle-foot conditions and may change during rehabilitation, limiting the generalizability of a single rocker profile. Third, the use of a pressure walkway, which is less accurate than force plates and is subject to hysteresis and per-sensor force thresholds, may have influenced the vertical GRF estimates. Moreover, full 3D kinetics and kinematics were not captured in this preliminary evaluation. Finally, the evaluation was restricted to level-ground walking at self-selected speed in a laboratory setting. The current prototype includes mediolateral contact plates that increase the limb’s effective width. While these are functional on flat surfaces, they may present stability challenges on uneven terrain, which were not examined in this preliminary study.

Future studies will involve larger cohorts, encompassing a variety of ankle and foot injuries and conditions. These studies will include expanded kinematic and kinetic measurements, and incorporate functional tasks commonly used in rehabilitation protocols, such as navigating slopes and stairs. Additionally, out-of-laboratory assessments will be conducted to evaluate the OLAFO’s robustness and clinical utility in real-world conditions, including irregular surfaces and uneven terrain. This will involve using body-worn sensors for quantitative gait assessment [85–87].

Further development efforts will aim to refine the device by: (1) integrating components for elastic energy storage and return with optimized stiffness [88, 89]; (2) enhancing the brace design for improved pressure distribution through iterative optimization guided by finite element analysis, similar to the methods employed in the development of transtibial prosthetic sockets [58, 59, 61]; (3) refining the design to accommodate slopes and uneven terrains, and (4) reducing the overall mass of the device.

## 5 Conclusion

The OLAFO prototype demonstrated that a custom load-bearing shank brace, adjustable side struts, and ROS-informed contact plates can together provide both complete offloading and graded partial loading while preserving relatively natural gait characteristics. In a healthy participant, the device achieved full plantar offloading, a nearly linear relationship between strut height and foot loading impulse, and only modest increases in contralateral stance duration and intact-limb GRFs compared with normal gait. While this initial evaluation focused on level-ground walking in a single participant to establish proof-of-concept, these findings support the feasibility of the OLAFO as a tool for prescribing and verifying partial weightbearing targets during rehabilitation. Future iterations will focus on optimizing the OLAFO’s profile for varied terrains and clinical safety, motivating further evaluation in patient populations and real-world clinical settings.

## Acknowledgments

We thank Etamar Bareket and Hadas Huber for their contributions to the OLAFO design. This research was supported by grants from the Israel Ministry of Science and Technology and by the Mark S. Kahn Family Fund, the J. Tal Fund, the Henri Gutwirth Fund, the Miriam and Aaron Gutwirth Fellowship, and the Neubauer Doctoral Fellowship. The funders had no role in study design, data collection and analysis, decision to publish, or preparation of the manuscript.

## Data Availability

Data can be made available to interested researchers upon request by email to the corresponding author.

